# Microstructural Properties of Human Brain Revealed by Fractional Anisotropy can Predict the After-effect of Intermittent Theta Burst Stimulation

**DOI:** 10.1101/2021.08.30.458153

**Authors:** Ikko Kimura, Hiroki Oishi, Masamichi J Hayashi, Kaoru Amano

**Affiliations:** Center for Information and Neural Networks (CiNet), Advanced ICT Research Institute, National Institute of Information and Communications Technology, Suita 565-0871, Japan; Graduate School of Frontier Biosciences, Osaka University, Suita 565-0871, Japan; Graduate School of Information Science and Technology, The University of Tokyo, Tokyo 113-8656, Japan

**Keywords:** DTI, inter-individual variability, fractional anisotropy, iTBS, rTMS

## Abstract

Intermittent theta burst stimulation (iTBS) delivered by transcranial magnetic stimulation (TMS) produces a long term potentiation (LTP)-like after-effect useful for investigations of cortical function and of potential therapeutic value. However, the iTBS-evoked after-effect over the primary motor cortex (M1) as measured by changes in motor evoked potential (MEP) amplitude exhibits a largely unexplained variability across individuals. Here, we present evidence that individual differences in white and grey matter microstructural properties revealed by fractional anisotropy (FA) predict the magnitude of the iTBS-induced after-effect over M1. The MEP amplitude change in the early phase (5–10 min) post-iTBS was associated with FA values in white matter tracts such as right superior longitudinal fasciculus and corpus callosum. In contrast, the MEP amplitude change in the late phase (15–30 min) post-iTBS was associated with FA in grey matter, primarily in right frontal cortex. These results suggest that the microstructural properties of regions connected directly or indirectly to the target region (M1) are crucial determinants of the iTBS after-effect. FA values indicative of these microstructural differences can predict the potential effectiveness of rTMS for both investigational use and clinical application.

## Introduction

Repetitive transcranial magnetic stimulation (rTMS) is widely used to modulate cortical excitability for experimental investigations and for the treatment of diseases such as major depression, movement disorders and chronic pain (Lefaucheur et al. 2020). Thus, it is of great experimental and clinical value to predict those subjects or patients most responsive prior to application. Intermittent theta burst stimulation (iTBS) is a rTMS protocol consisting of three pulses at 50 Hz repeated at 200-ms intervals (5 Hz) and delivered intermittently for 191 s (600 pulses in total) (Huang et al. 2005). This method is currently the focus of intensive preclinical and clinical investigation (Suppa et al. 2016; Rounis and Huang 2020) as this pattern can evoke a long term potentiation (LTP)-like after-effect in corticospinal excitability lasting for around 30 min when applied over primary motor cortex (M1) (Huang et al. 2005; López-Alonso et al. 2014), significantly longer than conventional rTMS protocols (e.g. continuous 5 Hz rTMS) using the same stimulation length.

A major concern when using iTBS, however, is the substantial inter-individual variability in the magnitude of this after-effect (Hamada et al. 2013; Hinder et al. 2014; López-Alonso et al. 2014; Leodori et al. 2021; Ozdemir et al. 2021). Thus, prior assessment of iTBS susceptibility would be useful for obtaining robust results in rTMS experiments (López-Alonso et al. 2014) and for identifying patients most likely to benefit from clinical application. Previous studies have found that age, genetic polymorphisms, time of day iTBS is delivered and hormone levels are associated with inter-individual variability in the after-effect (for review, see (Suppa et al. 2016)). However, these factors are not region specific, and underlying neuro-modulatory mechanisms critical for the iTBS-evoked after-effect are unknown.

Recently, Nettekoven and colleagues reported that the functional connectivity (FC) values between M1 and other cortical regions predicted the magnitude of the iTBS after-effect (Nettekoven et al. 2015). This result suggests that the strengths of neural connections with iTBS targets may also influence the inter-individual variability. However, given that FC fluctuates depending on the subject’s state of mind and alertness (Rosenberg et al. 2016), anatomical connectivity, which is independent of these factors, may be a more robust predictor. Fractional anisotropy (FA), a metric of diffusion-weighted magnetic resonance imaging (dMRI) that quantifies the anisotropy in directionality of water diffusion, is associated with the microstructural properties of neural tissue such as cell density and the orientation, diameter and myelination of axons (Le Bihan et al. 2001; Beaulieu 2002; Le Bihan 2003). Further, several studies have found that FA values can predict the ability to learn new motor skills (Tomassini et al. 2011; Schulz et al. 2015; Lehmann et al. 2019) and the recovery rate of motor function after stroke (Kumar et al. 2016; Puig et al. 2017; Soulard et al. 2020), suggesting that FA is associated with synaptic plasticity within motor-associated regions.

We speculated that the microstructural properties reflected by dMRI are associated with inter-individual differences in the after-effect of iTBS. To test this hypothesis, we examined whether regional FA values within the human brain, which reflect local microstructural properties of white and grey matter, are correlated with inter-individual variability in the iTBS-induced after-effect over M1. We report that FA values in certain white and grey matter regions predict the magnitude of the after-effect during the early and late phases, respectively, while other factors, such as sex and age, had little influence. These findings suggest that metrics derived from dMRI measurement, such as FA, are strong predictive indicators of regional responses to iTBS and possibly other rTMS protocols.

## Materials and Methods

The experiment required 2 days for completion by each participant. On day 1, MRI data were collected. On day 2, the after-effect of iTBS was assessed by measuring the amplitude of motor evoked potentials (MEPs) before (baseline) and after iTBS (Fig. 1a).

**Figure. 1.**
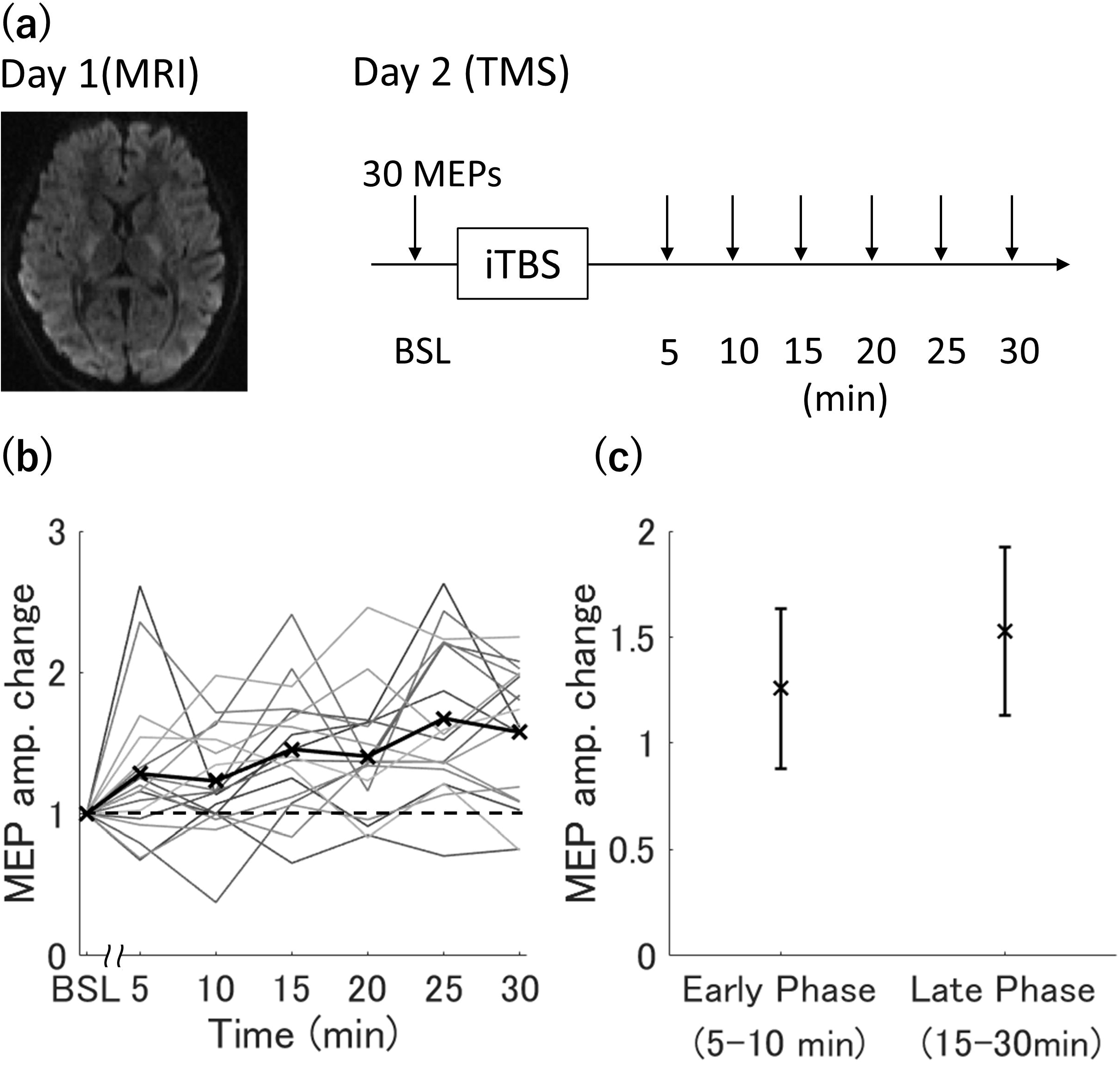
Overview of the experimental protocol for measuring changes in motor evoked potential (MEP) amplitude after intermittent theta burst stimulation (iTBS) over the primary motor cortex. (a) On day 1, anatomical MRI data were acquired. On day 2, a transcranial magnetic stimulation (TMS) experiment was performed, in which MEPs were measured before and after iTBS. (b) Individual time courses of MEP amplitude changes for all participants (n = 18). The horizontal axis indicates the time after iTBS, while the vertical axis indicates MEP amplitude normalised to baseline amplitude (BSL). Thick black line and cross indicate the mean change in MEP amplitude across individuals. (b) Changes in MEP amplitude during the early phase (5–10 min after iTBS) and the late phase (15–30 min after iTBS). The black cross indicates the mean change and the error bars indicate ± 1 standard deviation.

### Participants

Eighteen healthy adult volunteers (13 males and 5 females; age 20–24 years; mean ± standard deviations (SD), 21.7 ± 1.0 years) participated in this study. All subjects were right-handed according to the Edinburgh handedness inventory (Oldfield 1971) and reported no history of neuropsychiatric diseases. The experiments were approved by the institutional ethics and safety committees of the National Institute of Information and Communications Technology and were performed in accordance with the Declaration of Helsinki. All participants provided informed consent after a full explanation of study protocols and aims.

### Transcranial Magnetic Stimulation

Each iTBS session was performed in the afternoon, starting from around 1 pm or 3 pm, to mitigate known diurnal variation in response. To determine the optimal stimulator output for MEP measurement and iTBS, resting motor threshold (RMT) and active motor threshold (AMT) were defined for each participant prior to the iTBS session. The RMT was defined as the lowest intensity that evoked a MEP of at least 50 μV on 5 out of 10 trials in the contralateral first dorsal interosseous (FDI) muscle at rest (Rossini et al. 2015), while AMT was defined as the lowest intensity that evoked a MEP of at least 200 μV on 5 out of 10 trials in the contralateral FDI muscle during volitional contraction at ~10% of maximum (Rossini et al. 2015). Thirty MEPs from the left FDI muscle were recorded ~10 min before iTBS (baseline) and for up to 60 min after iTBS at 5-min intervals, with the stimulator output set to 120% of the RMT. Before iTBS, alertness of the participants was also assessed by the Stanford Sleepiness Scale (SSS) (Hoddes 1972). All MEPs for determination of RMT and the iTBS after-effect were evoked using a monophasic Magstim 200^2^ stimulator (Magstim, Whitland, UK) with a figure-of-eight 70-mm standard coil, while AMT was determined using a biphasic Magstim Rapid^2^ stimulator (Magstim, Whitland, UK) with a figure-of-eight 70-mm air film coil.

The iTBS was delivered over the right M1 using the same stimulator and coil as used for AMT measurements. The stimulation was targeted to the hotspot over the M1 evoking the strongest MEP in the right FDI muscle. Coil orientation was also optimised to elicit the largest MEPs, with the coil handle pointing backward and ~45° from the midline. The iTBS protocol was the same as that introduced by Huang et al. (Huang et al. 2005), consisting of 2-s trains of three pulses at 50 Hz repeated every 200 ms (5 Hz). Trains were repeated 20 times at an 8-s inter-train interval for 191 s (a total 600 pulses). The stimulation intensity was usually set to 80% of the AMT. However, when 80% AMT exceeded the system’s upper limit for TBS protocols (corresponding to 50% maximum stimulator output (MSO)), the stimulator output was set at this upper limit. The coil position was monitored and recorded using a Brainsight neuronavigation system (Rogue Research Inc, Montreal, Canada).

### Electromyography

Motor evoked potentials from the left FDI muscle were recorded as surface electromyogram (EMG) signals using pre-gelled Ag-AgCl electrodes, with the active electrode placed on the muscle belly and the reference electrode on the metacarpophalangeal joint of the left index finger. The MEP signals were amplified and recorded with 16 Hz to 470 Hz band-pass filtering and 3 kHz digitisation using Brainsight (Rogue Research Inc, Montreal, Canada).

### Motor evoked Potential Analysis

MEP amplitudes were measured using custom software and the Veta-Toolbox (Jackson and Greenhouse 2019) implemented in Matlab (MathWorks, Natick, Massachusetts). For three participants, we were unable to measure MEPs 35 min after iTBS due to coil overheating and hence only the time points up to 30 min post-iTBS were used for subsequent analysis. At each time point post-iTBS, MEPs larger than 2.5 SDs from the mean of 30 trials were rejected as outliers (Fried et al. 2017) and the remaining MEP amplitudes were averaged and divided by the mean baseline MEP amplitude to calculate the after-effect of iTBS. Since previous studies have reported bimodal changes in MEP amplitude after iTBS, with a trough at 12.5 min post-iTBS (Huang et al. 2005), we separated the time points into two phases, early (5 and 10 min after iTBS) and late (from 15 to 30 min after iTBS) and the change in amplitude at each time point within each phase relative to baseline was averaged for subsequent analysis. To validate this phase stratification, we performed hierarchical clustering analysis (Supp. 1). First, similarity measures were calculated between each time point as the Euclidean distance between vectors of MEP amplitude for all participants and then clustering analysis was conducted using the single linkage method (Murtagh and Contreras 2012).

### Image acquisition

Prior to the iTBS session, diffusion-weighted MRI (dMRI) data with b = 1000 s/mm^2^ and b = 2000 s/mm^2^ were collected from each participant using a Siemens Vida 3T scanner and 64-channel array head coil (Siemens, Erlangen, Germany). Both diffusion-weighted images were obtained using a multislice 2D single-shot spin-echo echo-planar sequence with the following parameters: voxel size = 2 × 2 × 2 mm, matrix size = 106 × 106 × 74, iPAT reduction factor = 2, Multiband Acceleration Factor = 3, phase-encoding direction = A-P, and TR = 5300 ms. The number of directions and TE differed between the two datasets, with number of directions = 30 and TE = 71 ms for the b = 1000 s/mm^2^ dataset and number of directions = 60 and TE = 86 ms for the b = 2000 s/mm^2^ dataset. Eight non-diffusion-weighted (b = 0 s/mm^2^) images were also acquired to minimise EPI distortion, five images with the same TE as in the b = 1000 s/mm^2^ dataset with three images reversed phase-encoding directions (i.e. P-A), and three images with the same TE and phase-encoding directions (i.e. A-P) as in the b = 2000 s/mm^2^ dataset. Total acquisition time for dMRI was around 10 min for each participant.

For neuronavigation of TMS coil position and surface-based analysis, a T1-weighted MP-RAGE image was also obtained for each participant (voxel size = 1 × 1 × 1 mm, TE = 2.48 ms, TR = 1900 ms, flip angle = 9°).

### Image Analysis

The dMRI data were preprocessed using tools from the FMRIB software library (FSL 6.0.1, https://fsl.fmrib.ox.ac.uk/fsl). The FSL topup tool was used to correct for EPI distortions due to inhomogeneity in the magnetic field (Andersson et al. 2003) and the eddy tool was used to correct for eddy current (Andersson and Sotiropoulos 2016) with outlier replacement and slice-volume correction (Andersson et al. 2016). Non-brain tissue was removed using Brain Extraction Tool (Smith 2002). FA was calculated for each b = 1000 s/mm^2^ and b = 2000 s/mm^2^ dataset using the DTIFIT application of FSL, and averaged across individual datasets to obtain individual FA maps.

To determine whether the difference in FA reflects the density of neural cells and/or the dispersion of neuronal fibre orientation (Zhang et al. 2012), the neural density index (NDI) and orientation dispersion index (ODI) were calculated using the NODDI toolbox v1.0.3 (www.nitrc.org/projects/noddi_toolbox). Before fitting the NODDI model, diffusion-weighted images at each b value were divided by the mean non-diffusion-weighted image obtained with the same TE value to merge each dataset (Owen et al. 2014; Chang et al. 2015; Palacios et al. 2020). The fitting was performed using the default settings for white matter (WM), while the intrinsic free diffusivity parameter was changed to 1.1 × 10^-3^ mm^2^/s for grey matter (GM) (Fukutomi et al. 2018, 2019; Guerrero et al. 2019).

The Tract-Based Spatial Statistics (TBSS) tool of FSL (Smith et al. 2006) was used for whole-brain voxel-wise analysis of WM. First, the FA map for each participant was non-linearly registered to 1×1×1 mm^3^ MNI152 space (McConnell Brain Imaging Centre, Montreal Neurological Institute). From the mean FA image across participants, a common skeleton was extracted to represent the main WM structure. This skeleton was thresholded at FA > 0.2 (Default) and FA data warped to MNI152 space were then projected onto this skeleton. The NDI and ODI were also projected onto the skeleton using the same warp used for FA.

For surfaced-based analysis, FreeSurfer (Version 6.0.0, https://surfer.nmr.mgh.harvard.edu/) was first used to obtain individual cortical surfaces (Fischl et al. 1999). After removing non-brain tissues, the structural brain image was normalised into Talairach space and the intensity of each image was normalised. The normalised brain image was then segmented into GM, WM and cerebral spinal fluid (CSF). The GM–WM boundary (WM surface) and GM–CSF boundary (pial surface) were used for surface reconstruction. Utilising the folding pattern, the surface was registered to the standard surface space (fsaverage). All dMRI-derived GM metrics were sampled from the mid-points of white and pial surfaces as the partial-volume effect is less likely to impact the results (McNab et al. 2013). The sampled data was warped to fsaverage space, and smoothed with a Gaussian Kernel of 10-mm FWHM across the cortical surface according to a previous study using FA for surface-based analysis (Stock et al. 2020).

### Statistical analysis

The FSL randomise tool was used to test the statistical significance of associations between each phase of MEP amplitude change and the TBSS data. For the surface-based analysis, the FSL Permutation Analysis of Linear Models tool was used to test these associations in GM. In both WM and GM, we performed 5000 permutation tests and employed Threshold-Free Cluster Enhancement to control for multiple testing (Smith and Nichols 2009). To specify locations significantly correlated with MEP amplitude change, significantly associated WM and GM voxel clusters were labelled using the Johns Hopkins University white-mater tractography atlas (https://fsl.fmrib.ox.ac.uk/fsl/fslwiki/Atlases) and Desikan-Killiany atlas (Desikan et al. 2006), respectively. To distinguish the microstructural properties contributing to inter-individual variability in FA, we calculated Pearson’s correlation coefficients between mean FA and mean NDI or ODI in each cluster and tested the significance by 5000 permutation tests (Supp. 5). In all analysis, Bonferroni correction was used to adjust for possible spurious findings due to multiple testing.

We also calculated Pearson’s correlation coefficients between all measured continuous variables and MEP amplitude change within each phase, while two-tailed unpaired t-tests were performed to assess how MEP amplitude change is affected by sex and iTBS intensity (80% of AMT or 50% of MSO). A P < 0.05 was considered significant for all tests. These analyses were performed using JASP (ver. 0.13.1 for Windows, https://jasp-stats.org/)

## Results

All participants completed both MRI scans and iTBS experiments, and no adverse events occurred during these procedures. Plotting the individual time courses of MEP amplitude changes post-iTBS relative to baseline revealed substantial inter-individual variation (Fig. 1b), in accord with previous reports (Hamada et al. 2013; López-Alonso et al. 2014). Consistent with the previous study showing the two phases of MEP amplitude change after iTBS (Huang et al. 2005), our hierarchical clustering analysis revealed the similar separation in the time course (early phase; 5–10 min after iTBS, late phase; 15–30 min after iTBS) (Supp. 1). Therefore, the relationships between post-iTBS MEP amplitude and brain microstructural properties were analysed separately for the early and late phases.

We first tested the associations between regional FA and MEP amplitude change. During the early phase, MEP amplitude change was significantly and negatively correlated with FA of WM tracts anatomically connected to M1 (Fig. 2a; for the spatial maps in all slices, see Supp. Fig. 2), including right superior longitudinal fasciculus, right posterior corona radiata, right internal capsule and right corticospinal tract. This voxel cluster is defined as WM 1 and other significant clusters as WM 2–10 in Figure 2a. There were also significant negative correlations between MEP amplitude change and FA in bilateral corpus callosum (WM 2 and 3), right forceps minor (WM 4 and 5), left anterior limb of the internal capsule (WM 6), left retrolenticular part of the internal capsule (WM 7), left posterior limb of the internal capsule (WM 8) and right uncinate fasciculus (WM 9 and 10). Conversely, regional FA in the GM was not correlated with the early-phase MEP amplitude change.

**Figure. 2.**
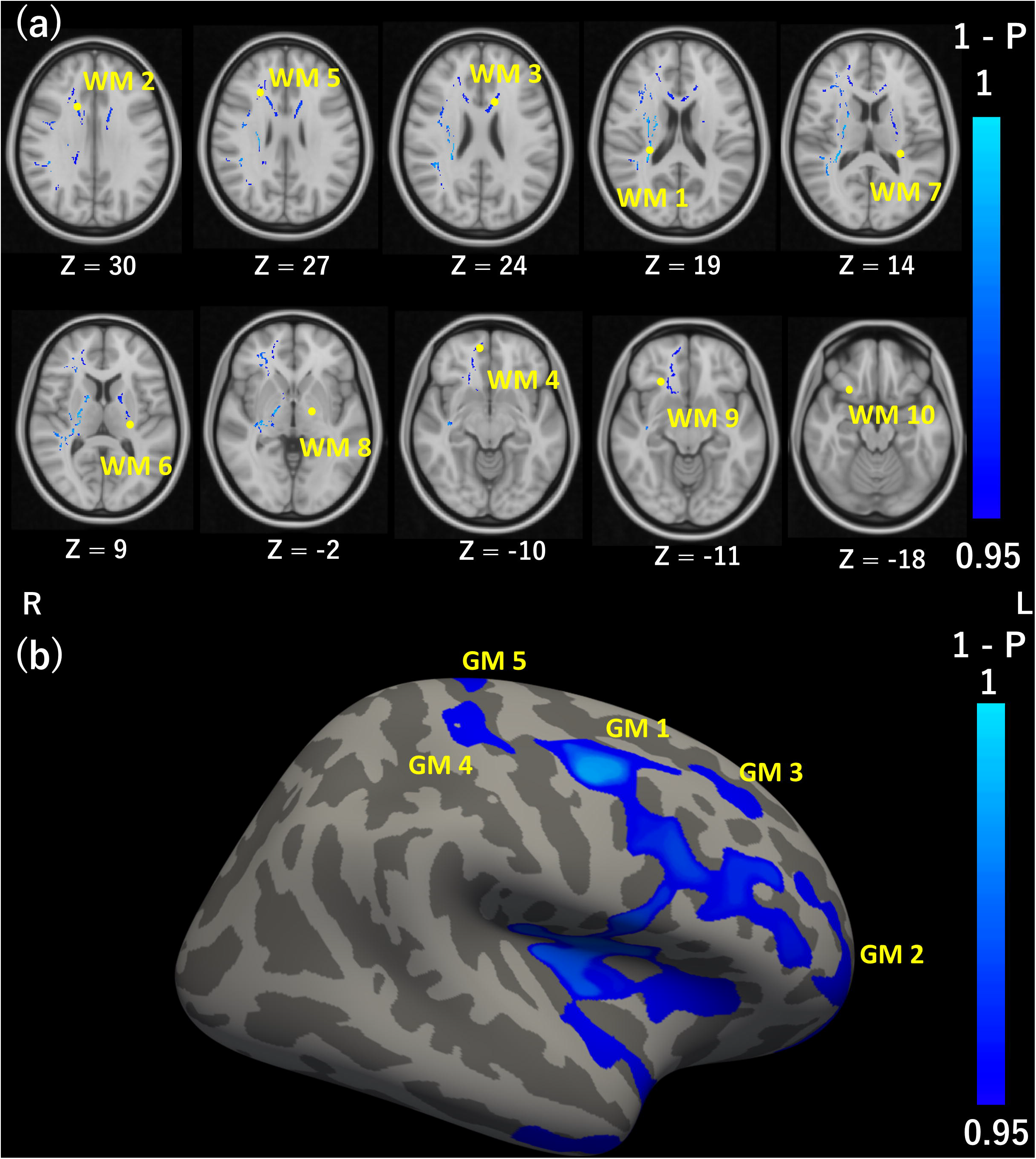
Correlations between regional fractional anisotropy (FA) and motor evoked potential (MEP) amplitude changes during early and late phases. (a) Tract-Based Spatial Statistics (TBSS) analysis of significant correlations between FA in white matter (WM) and early-phase MEP amplitude change. Clusters in blue represent voxels showing significant negative correlations with FA. The numbers (WM 1–10) correspond to individual voxel clusters. Yellow dots indicate the lowest T-values (strongest correlations) in each cluster. The axial slices are displayed according to radiological convention (left on the picture is right in the brain) with the range in MNI coordinates from z = 30 (top left) to z = −18 at bottom right. (b) Surface-based analysis of significant correlations between FA in grey matter (GM) and late-phase MEP amplitude change. Clusters in blue represent voxels showing significant negative correlations with FA. The numbers (GM 1–5) correspond to the individual voxel clusters.

In contrast to the early phase of MEP amplitude change, the late-phase change was negatively correlated with FA exclusively in GM regions (Fig. 2b), including the right caudal middle frontal region, right pars opercularis, right insula, right superior temporal and right middle temporal regions (GM 1). Negative correlations were also found between late-phase MEP amplitude change and FA in the anterior part of the right rostral middle frontal region (GM 2), dorsal part of the right rostral middle frontal region (GM 3), right postcentral gyrus (GM 4) and right precentral and postcentral gyrus (GM 5). There were no significant correlations between the late-phase MEP amplitude change and FA in the WM.

We also tested whether inter-individual differences in MEP amplitude change during the early and late phases were associated with other collected measurements (Table 1). Age, handedness, and interval between the MRI scan and iTBS session were not significantly correlated with either early- or late-phase MEP amplitude change (P > 0.05). Similarly, stimulus intensity for MEP induction, iTBS intensity, Stanford sleepiness scale score and mean baseline MEP amplitude were not significantly correlated with MEP amplitude change (P > 0.05). Unpaired t-test also revealed no significant differences in MEP amplitude change between sexes (early phase, t = 0.77, P = 0.45; late phase, t = 0.26, P = 0.80) or between subjects receiving iTBS at 80% of AMT or 50% of MSO (early phase, t = 1.27, P = 0.33; late phase, t = −0.23, P = 0.82).

**Table 1.**
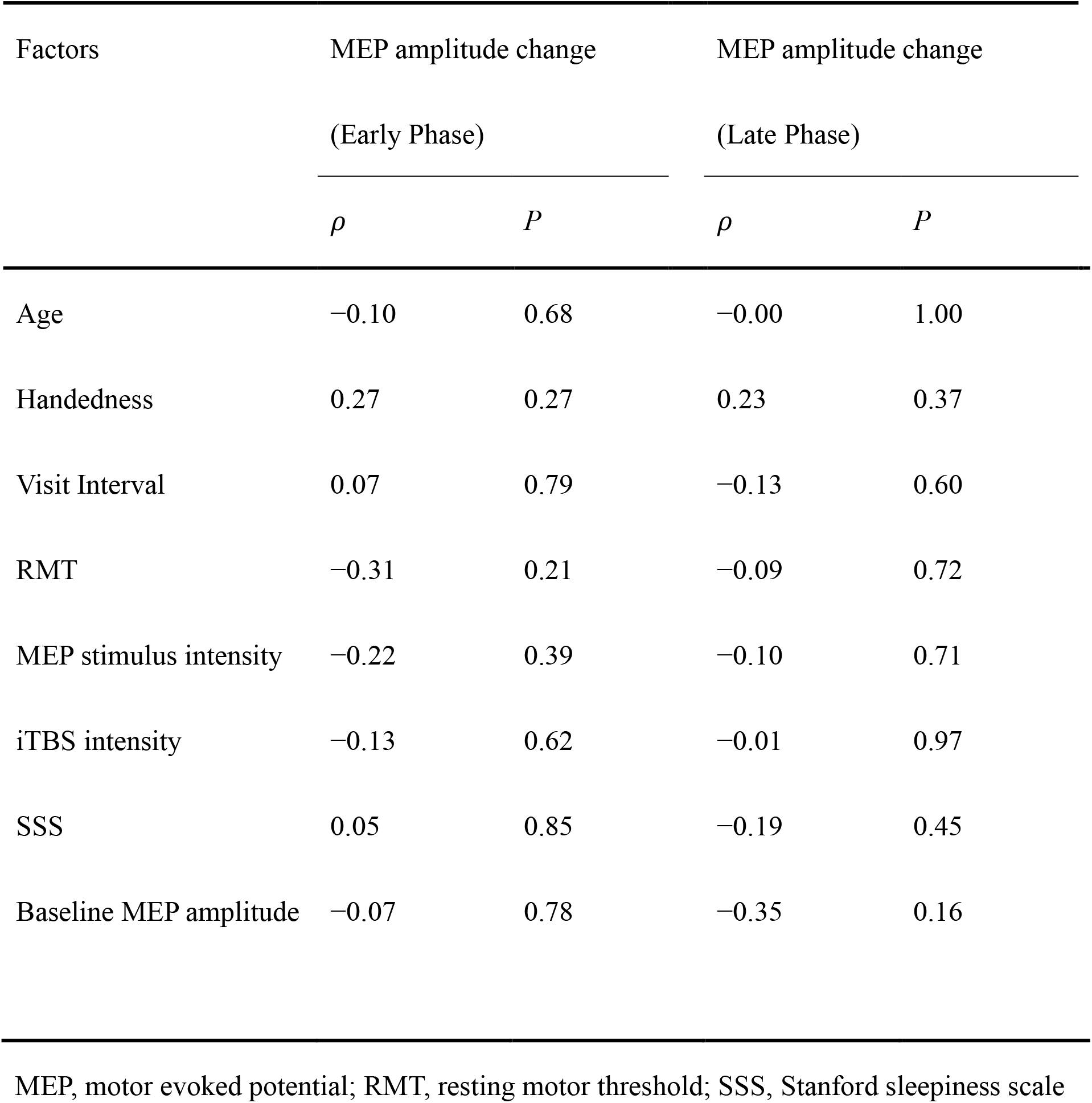
Correlations between MEP amplitude change and other measured factors

| Factors | MEP amplitude change |  | MEP amplitude change |  |
| --- | --- | --- | --- | --- |
|  | (Early Phase) |  | (Late Phase) |  |
| | $\rho$ | $P$ | $\rho$ | $P$ |
| Age | −0.10 | 0.68 | −0.00 | 1.00 |
| Handedness | 0.27 | 0.27 | 0.23 | 0.37 |
| Visit Interval | 0.07 | 0.79 | −0.13 | 0.60 |
| RMT | −0.31 | 0.21 | −0.09 | 0.72 |
| MEP stimulus intensity | −0.22 | 0.39 | −0.10 | 0.71 |
| iTBS intensity | −0.13 | 0.62 | −0.01 | 0.97 |
| SSS | 0.05 | 0.85 | −0.19 | 0.45 |
| Baseline MEP amplitude | −0.07 | 0.78 | −0.35 | 0.16 |

MEP, motor evoked potential; RMT, resting motor threshold; SSS, Stanford sleepiness scale

Finally, to identify factors contributing to FA values in each voxel cluster associated with MEP amplitude change, we investigated the correlations between mean FA and both mean NDI and mean ODI, which respectively reflect neural cell density and the dispersion of fibre orientation within a voxel (Zhang et al. 2012). The FA was positively correlated with NDI in WM 1 (right superior longitudinal fasciculus and right corticospinal tract), WM 4 and 5 (right forceps minor) and WM 6 (left anterior limb of internal capsule) (Supp. 3a), while FA was negatively correlated with ODI in all WM clusters (Supp. 3b). The FA values of GM 1–3 (right frontal regions, right insula and right temporal regions) were also negatively correlated with ODI (Supp. 4a), while FA values of GM 4–5 (the right postcentral gyrus) were positively correlated with NDI (Supp. 4b).

## Discussion

In this study, we demonstrated multiple significant associations between MEP amplitude change after iTBS over M1 (the iTBS-induced after-effect) and the microstructural properties of GM and WM regions associated with M1. Further, no other measured factors showed significant associations. Thus, individual variation in these microstructural properties can explain, at least in part, the known individual variation in iTBS after-effect, thereby providing a potential method to predict responsive individuals prior to neuroscientific investigations and possibly iTBS-based therapy.

The early-phase MEP amplitude change was negatively correlated with regional FA values in WM tracts including right superior longitudinal fasciculus, corpus callosum, right forceps minor, left internal capsule and right uncinate fasciculus. In contrast, the late-phase MEP amplitude change was negatively correlated with regional FA in GM areas such as right frontal cortex, right insula, right temporal cortex and right postcentral gyrus. Collectively, these findings might suggest that distinct mechanisms underlie the early and late phases of the iTBS-evoked after-effect.

### Negative correlations of regional WM FA with early-phase MEP amplitude change

The early-phase MEP amplitude change was negatively correlated with regional FA values in WM tracts related to motor function, including the right corticospinal tract (WM 1), right superior longitudinal fasciculus (WM 1) and corpus callosum (WM 2 and 3), which are functionally and anatomically connected to M1 (Catani and de Schotten 2012). The corticospinal tract sends output from M1 to the contralateral spinal cord and ultimately to distal muscles (Lemon 2008) and output strength (number of motor neurons recruited) determines MEP amplitude (Bestmann and Krakauer 2015). The superior longitudinal fasciculus is the main intrahemispheric tract connecting frontal areas (e.g. premotor cortex, dorsolateral prefrontal cortex (DLPFC), and M1) and parietal areas (e.g. angular gyrus and supramarginal gyrus), and is crucial for motor planning, motor imagery, and visuo-motor tasks (Nakajima et al. 2020). The corpus callosum connects the bilateral M1 (Hofer and Frahm 2006) and mediates both interhemispheric inhibition (Ferbert et al. 1992) and interhemispheric facilitation (Hanajima et al. 2001).

The FA values within a part of the corpus callosum that connects bilateral frontal regions (WM 2 and 3), right forceps minor (WM 4, 5, 9, and 10), and left internal capsule (WM 6-8) were also negatively correlated with MEP amplitude change. Given that these tracts are not directly involved in motor function, the negative correlations may be mediated by factors affecting general WM functional properties, such as the gene polymorphisms of the brain derived neurotrophic factor, which are known to influence both the microstructural properties of WM (Chiang et al. 2011) and the iTBS after-effect (Cheeran et al. 2008; Antal et al. 2010).

Further examination of the associations between regional FA values in WM and both NDI and ODI, which respectively reflect the density of neural fibres and the dispersion of fibre orientation within a voxel (Zhang et al. 2012), provided clues to the nature of these microstructural differences underlying individual variation in iTBS-evoked MEP change. A larger MEP change was associated with lower FA in multiple tracts, and in several of these tracts (including right superior longitudinal fasciculus and right corticospinal tract of WM 1, right forceps minor in WM 4 and 5, and left anterior limb of internal capsule in WM 6), FA was negatively correlated with NDI (Supp. 3a), while FA and ODI were negatively correlated in all clusters (Supp. 3b). Therefore, lower FA values in WM 1, 4, 5, 6, and 10 are associated with larger MEP amplitude change, less consistent fibre orientation (higher dispersion), and a smaller neural fraction, possibly reflecting smaller diameter fibres or lower myelination. Alternatively, lower FA values in the other WM clusters may reflect only less consistent fibre orientation.

### Negative correlations of regional grey matter FA with late-phase MEP amplitude change

In contrast to the early phase, the late-phase MEP amplitude change was negatively correlated with regional grey matter FA, primarily in the right frontal cortex (GM 1–3). These regions include right premotor cortex (GM 1), right DLPFC (GM 1), and right anterior prefrontal cortex (aPFC) (GM 2), all of which are implicated in motor function. The premotor cortex is crucial for motor planning and sends information encoding these plans to M1 for execution (Hoshi and Tanji 2007), while the DLPFC integrates inputs from multiple sensory modalities to decide on the action to take (Yarrow et al. 2009) and the aPFC is important for motor response inhibition such as in go/no-go tasks (Boecker et al. 2007; Wriessnegger et al. 2012). Both the premotor cortex (Civardi et al. 2001; Koch et al. 2007; Bäumer et al. 2009; Groppa et al. 2012) and DLPFC (Hasan et al. 2013; Cao et al. 2018) modulate the activity of ipsilateral M1. Furthermore, premotor cortex was shown to modulate the plasticity of ipsilateral M1 (Huang et al. 2018).

The late-phase MEP amplitude change was also negatively correlated with FA in the right postcentral gyrus (GM 4 and 5) and right insula (GM 1). The postcentral gyrus provides sensory feedback to M1 (Kaelin-Lang et al. 2002) and M1 modulates activity of the postcentral gyrus (Katayama and Rothwell 2007), suggesting reciprocal functional connections. The insula is a part of the saliency network and facilitates motor responses indirectly via the anterior cingulate cortex (Menon and Uddin 2010).

The FA values of GM 1–3 (right frontal regions, right insula, and right temporal regions) were also negatively correlated with ODI (Supp. 4a), indicating that the greater MEP amplitude change associated with lower FA may reflect more complex dendritic branching (Zhang et al. 2012). By contrast, the FA values of GM 4–5 (the right postcentral gyrus) were positively correlated with NDI (Supp. 4b), suggesting that a larger late-phase after-effect may be facilitated by a lower density of apical dendrites (Zhang et al. 2012; Ball et al. 2013).

### Microstructural properties may influence the after-effect of iTBS

Several studies have suggested that distinct mechanisms underlie the early-phase and late-phase MEP amplitude changes following iTBS. In animals, Hoppenrath and Funke found that pyramidal neurons and inhibitory interneurons were facilitated 10 to 20 min after iTBS (i.e. during the early phase), while inhibitory interneurons were suppressed 20 to 40 min after iTBS (Hoppenrath and Funke 2013). Thus, the early phase may reflect enhanced excitatory transmission by projection neurons while the late phase reflects disinhibition. However, these findings do not explain why the FA values of non-stimulated regions were negatively correlated with MEP amplitude change. Thus, future animal studies are needed to investigate how the microstructural properties of regions directly or indirectly connected to M1 influence local excitability.

Surprisingly, the FA in M1 was not predictive of the MEP amplitude change, possibly because FA in M1 is not a sensitive measure of dendritic properties, which are major factors contributing to the after-effect of iTBS as discussed above. In M1, the number of neural cells is higher in layer 5 than layer 1, whereas most dendrites are located in layer 1 (von Economo 2008). The properties of layer 1 dendrites may not be well reflected by FA values in M1, and so cannot predict the iTBS after-effect. Future neuroimaging studies with higher spatial resolution are needed to determine whether the microstructural properties of GM in stimulated regions predict the after-effect magnitude following iTBS.

### Limitations

Potential limitations of this study include the use of FA to evaluate the microstructural properties of GM, as FA may not be as sensitive an indicator of GM properties due to the relatively large partial-volume effect (Aggarwal et al. 2015). To resolve this issue, we analysed FA only in the middle part of the grey matter (Stock et al. 2020) or along the major structures of the WM (Smith et al. 2006), which we believe helped to minimise the impact of the partial-volume effect. Furthermore, previous dMRI studies with higher spatial resolution have demonstrated the validity of FA for quantifying and distinguishing the structures of cortical layers (McNab et al. 2013; Aggarwal et al. 2015). Nonetheless, we cannot rule out the possibility that the analysed regions are affected by partial-volume effects from surrounding areas. Future neuroimaging studies with higher spatial resolution are needed to clarify how microstructural properties in regions associated with M1 influence the after-effect following iTBS.

## Conclusions

The microstructural properties of certain WM regions are negatively associated with the magnitude of the early-phase iTBS-evoked after-effect, while the microstructural properties of certain grey matter regions are negatively associated with the magnitude of the late-phase iTBS-evoked after-effect. These results suggest that FA measured by dMRI can be a powerful tool for prediction of experimental and therapeutic iTBS responses.

## Acknowledgement

We thank Ian Greenhouse for advice on EMG data analysis and Tomoya Kawashima for critical comments on the manuscript. This work was supported by the Japan Society for the Promotion of Science (Grants-in-Aid for Scientific Research JP18H05523 to K.A., JP18H01101 to M.J.H., Grant-in-Aid for Scientific Research on Innovative Areas JP19H05313 to M.J.H., and Grants-in-Aid for JSPS Research Fellow JP20J11101 to H.O.) and Japan Science and Technology Agency (PRESTO JPMJPR17J1 to K.A. and JST PRESTO JPMJPR19J8 to M.J.H).

