## Supplementary Figure 1, Supplementary Figure 2, Supplementary Figure 3, Supplementary Figure 4 for "Microstructural Properties of Human Brain Revealed by Fractional Anisotropy can Predict the After-effect of Intermittent Theta Burst Stimulation"

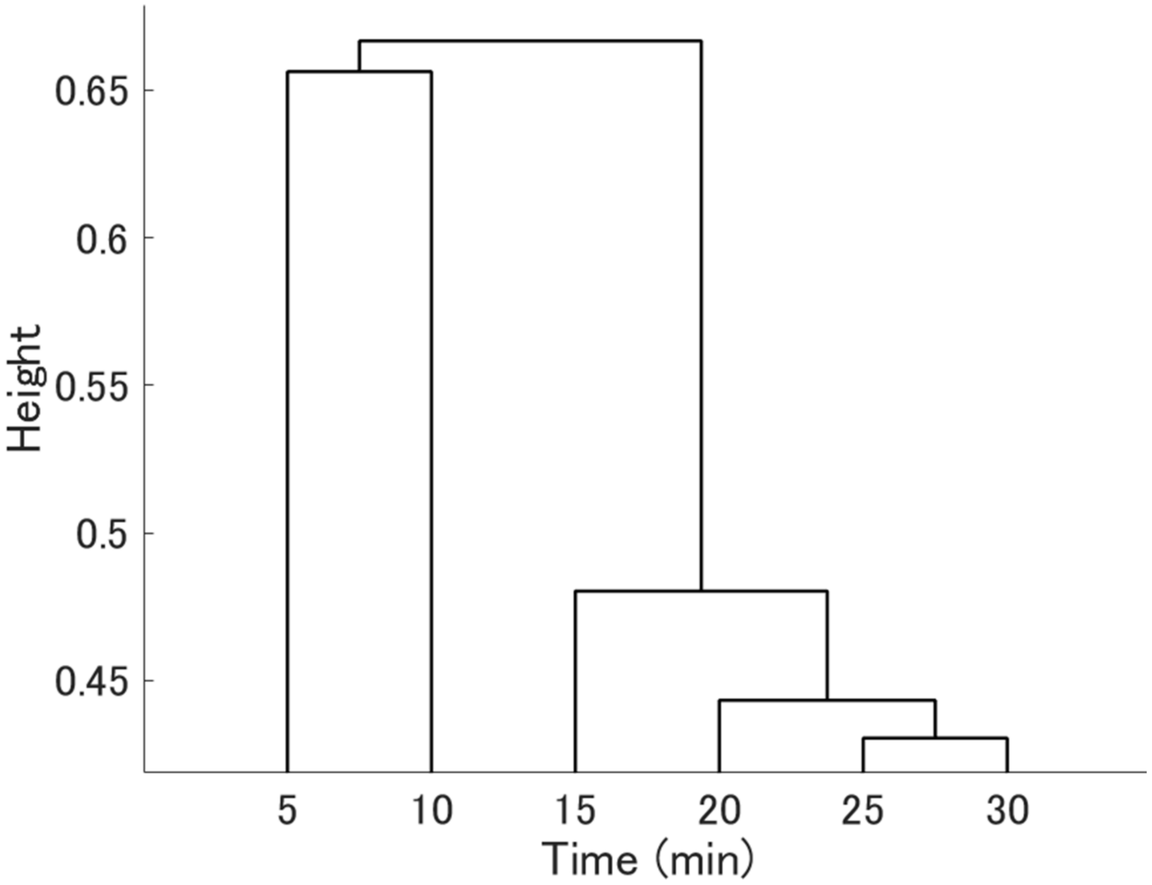


**Supplementary Figure 1.** Dendrogram of hierarchical clustering across the time course of motor evoked potential (MEP) amplitude change confirming distinct early and late phases. The vertical axis indicates the time after iTBS and the horizontal axis indicates the Euclidean distance.

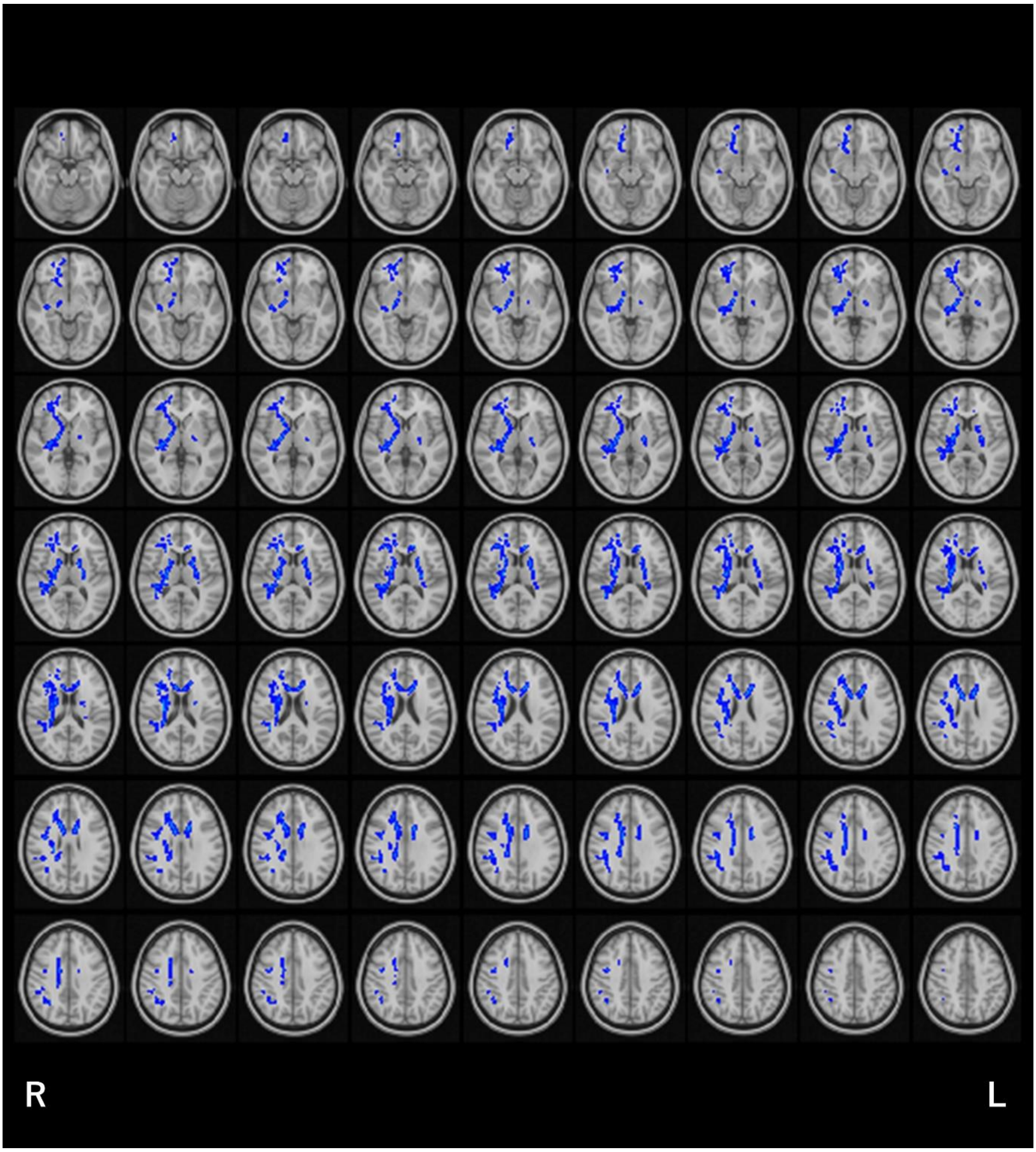


**Supplementary Figure 2.** Spatial map of correlations between fractional anisotropy (FA) and early-phase motor evoked potential (MEP) amplitude change in all slices (z = −18–44) rendered on the standard MNI template. Blue voxels represent regions in which FA was negatively correlated with early-phase MEP amplitude change. The significant voxels are filled using the FSL tbss_fill application to aid visualisation.

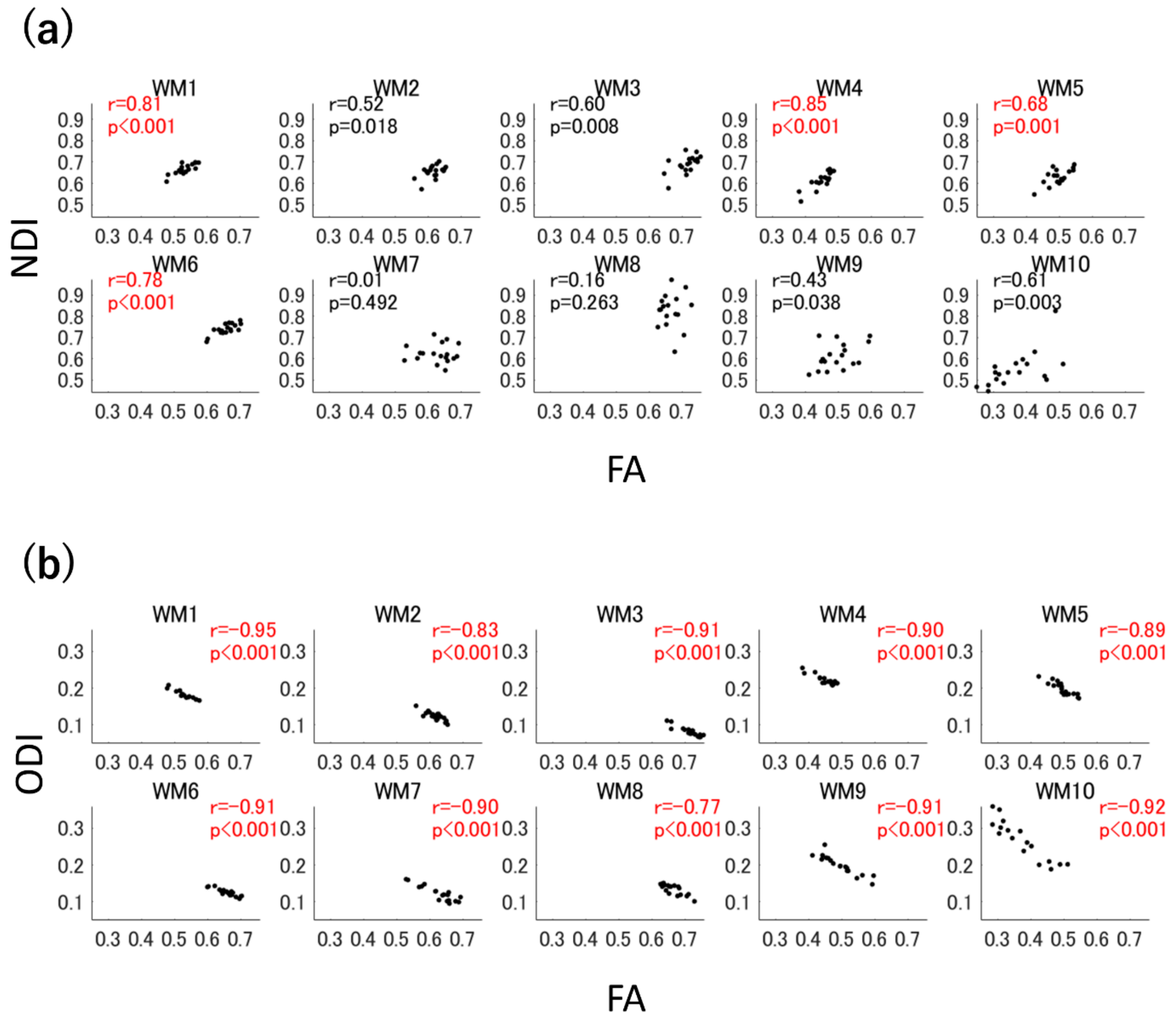


**Supplementary Figure 3.** (a) Scatter plots of the mean neural density index (NDI) versus the mean fractional anisotropy (FA) in each cluster showing significant negative correlations with early-phase motor evoked potential (MEP) amplitude change. Cluster numbers (WM 1–10) correspond to the labels assigned in Figure 1. The horizontal axis shows the mean NDI in each cluster, while the vertical axis shows the mean FA in each cluster. (b) Scatter plots of mean orientation dispersion index (ODI) versus mean FA in each cluster showing significant negative correlations with early-phase MEP amplitude change. The horizontal axis shows the mean ODI in each cluster, while the vertical axis shows the mean FA in each cluster. The text in red indicates that Pearson's correlation coefficient was significant after Bonferroni correction (i.e. $P < 0.05/20 = 0.0025$).

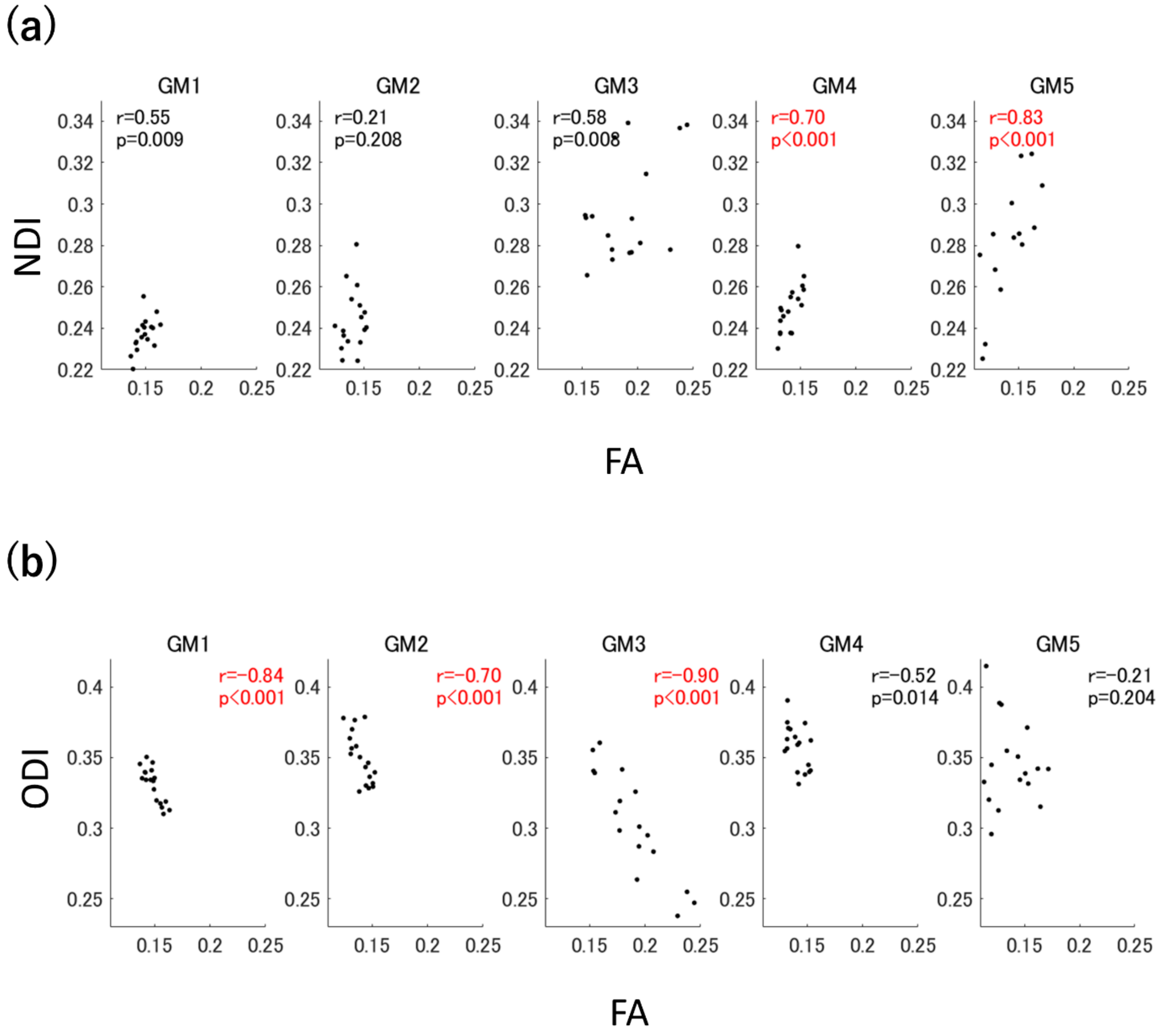


**Supplementary Figure 4.** (a) Scatter plots of the mean neural density index (NDI) versus the mean fractional anisotropy (FA) in each cluster that showed significant negative correlations with the late-phase motor evoked potential (MEP) amplitude change. Cluster numbers (GM 1–5) correspond to the labels assigned in Figure 1. The horizontal axis shows the mean NDI in each cluster, while the vertical axis shows the mean FA in each cluster. (b) Scatter plots of mean orientation dispersion index (ODI) versus mean FA in each cluster showing significant negative correlations with late-phase MEP amplitude change. The horizontal axis shows the mean ODI in each cluster, while the vertical axis shows the mean FA in each cluster. The text in red indicates that Pearson's correlation coefficient was significant after Bonferroni corrections (i.e. $P < 0.05 / 10 = 0.005$).
